# Sampling Neuron Morphologies

**DOI:** 10.1101/248385

**Authors:** Roozbeh Farhoodi, Konrad Paul Kording

## Abstract

The intricate morphology of neurons has fascinated since the dawn of neuroscience, and yet, it is hard to synthesize them. Current algorithms typically define a growth process with parameters that allow matching aspects of the morphologies. However, such algorithmic growth processes are far simpler than the biological ones. What is needed is an algorithm that, given a database of morphologies, produces more of those. Here, we introduce a generator for neuron morphologies that is based on a statistical sampling process. Our Reversible Jump Markov chain Monte Carlo (RJMCMC) method starts with a trivial neuron and iteratively perturbs the morphology bringing the features close to those of the database. By quantifying the statistics of the generated neurons, we find that it outperforms growth-based models for many features. Good generative models for neuron morphologies promise to be important both for neural simulations and for morphology reconstructions from imaging data.

## 1 Introduction

The morphology of neurons is beautiful and exhibits many regularities [1]. We can see the morphology as being the result of a process that optimizes aspects of wiring cost [2] [3]. Alternatively, we can see morphology as being the result of a growth process [4]. There are also multiple other rule-like aspects. For example, Pierrets rule tells that there is a correlation between the neuron size and length of its axonal size[1] and Larkman’s rule tells us that diameter of segments in the neuron are reversely correlated with their length [5]. There are also important environmental influences on cellular morphologies [6] suggesting that neither simple optimization nor simple growth-rule approaches can be sufficient. There are many mechanisms at play in the process that generates neuron morphologies suggesting that a meaningful generator for neuron morphologies needs to be multifaceted.

The morphology of neurons is important as it affects the way neurons compute. Different parts of the dendritic tree can produce different kinds of signals. For example, the apical dendrite of layer 5 pyramidal neurons can produced Calcium spikes [7]. Similarly basal dendrites are often able to produce NMDA spikes [8]. Moreover, the travel of signals along the dendrites changes the signal transmission from a synapse to the soma [9]. The morphology of neurons is important for any kind of precise neuron simulation [10, 11]. As such, morphologies are an important aspect of neuroscience.

Driven by recent interests in brain simulation as well as neuronal reconstruction, there is renewed interest in generative models for morphologies. Scientists may want to simulate more neurons than morphologies characterized by anatomists. This produces a need for generating morphologies that are like those that have been characterized[12]. Moreover, when trying to reconstruct neurons from 3D imaging data [13][14] [15, 16] we could also benefit from good generative models as a deviation may indicate a reconstruction mistake. Both of these fields could benefit considerably from the existence of good generative models for neuron morphologies.

Today’s generators use multiple different intuitions [17]. One set of approaches uses a simple growth process. For example, they may start at the soma and, at every potential branching point sample from statistical descriptors[18] an idea which is used in popular tools [19, 20, 21]. In that approach, all decisions about growth are strictly local at each subsequent branching point. A second set of approaches tries to follow the growth process and utilizes knowledge about the simultaneously developing segments [22, 23, 24]. These approaches effectively bootstrap off the idea of optimization [2]; however decisions are still unaffected by geometry. A third set of approaches includes knowledge of the geometry, e.g. by simulating neurotrophic particles [25]. This approach allows fast generation but ignores all aspects that are directly related to the growths process. All these approaches are based on the idea of producing neurons based on insights into the way neuron generation works, are conceptually beautiful, and successfully describe important aspects of morphologies.

Neuron morphologies have many characterized properties. They have certain shapes, certain distributions of branch angles, densities etc. Hence, it is hard to know how well a given generator characterizes real neurons. Each method may be good, or even provably optimal, at generating certain features. But if we acknowledge that the real generative process is more complicated than these generators we are faced with the problem of how we could generate neurons that obey many of the aspects of real neuron morphologies. We want to have a generator that can match many features from the dataset.

If we have a feature set that allows us to ask how probable a hypothesized morphology is based on a model derived from a database of real morphologies then we could generate neurons through a sampling process. Such an approach could try lots of neuron morphologies and basically find out which ones are more like those of real neurons and thus probable. In [26] such a process is used for generating textures. Generalizing this idea to morphologies is hard because the topological structure of the ambient space is nested. In fact, Markov chain Monte Carlo (MCMC) is a set of generally applicable methods that can be used to sample if we have a function that approximates the probability. In the case of morphologies that could be the probability of the morphology induced from a model fit to the feature set. What makes the problem complicated is the fact that different morphologies are not just different in the settings of parameters but about their number. In such cases Reversible jump MCMC is appropriate for sampling [27]. Such an approach would come with the promise of satisfying any number of features of the neuron morphology.

Here we present an approach based on Markov chain Monte Carlo methods for generating morphologies. By iterating changes, the chain becomes closer and closer to the generative distribution until it samples from a meaningful distribution. The advantage of our approach is that it generates morphologies based on a dataset of neurons, compared to other generative models which we can only optimize a small number of generator parameters. In this setup by adding more features or generally by making the generative model better, the resulting morphologies gradually become more realistic. We study the convergence and mixing time of the method and compare it with a recent model for generating morphologies.

## 2 Results

Morphologies are beautiful and important, raising the question of how they can be synthesized. We thus introduce an approach that generates new samples for a class of neurons by analyzing the morphologies in an existing database. We first introduce a rich set of features for quantifying the morphologies of a neuron class. Although the values of the features vary across the neurons, we can extract their salient statistics and join all features into one vector. This, along with a naïve Bayes assumption, allows us to build a probabilistic model that represents a neuron class. Having probability distribution over the space of possible neurons enables us to use popular methods for extracting samples such as Reversible Jump Markov Chain Monte Carlo (RJMCMC). We will thus be able to construct morphologies that are statistically matched to those of the database.

### 2.1 Generative model

The similarity of neurons can be readily defined in a relevant feature space. Such a feature set could be learned from a large database [28], but here we use a hand-engineered feature set. Relevant features may be a real number, e.g. the total length of neuritis. They may be counts in a histogram, e.g. all the angles at the branching nodes within a certain interval (figure 1.a). They may also be a vector, e.g. the density of neuritis in cylindrical coordinates. While some real valued features reveal the global structure of a neuron, histograms and densities can describe the arborizations and branching patterns. By concatenating all the features, we represent each morphology by a long feature vector (figure 1.b). The difference in the features and hence distance of feature vectors can be used to compare two neurons.

**Figure 1:**
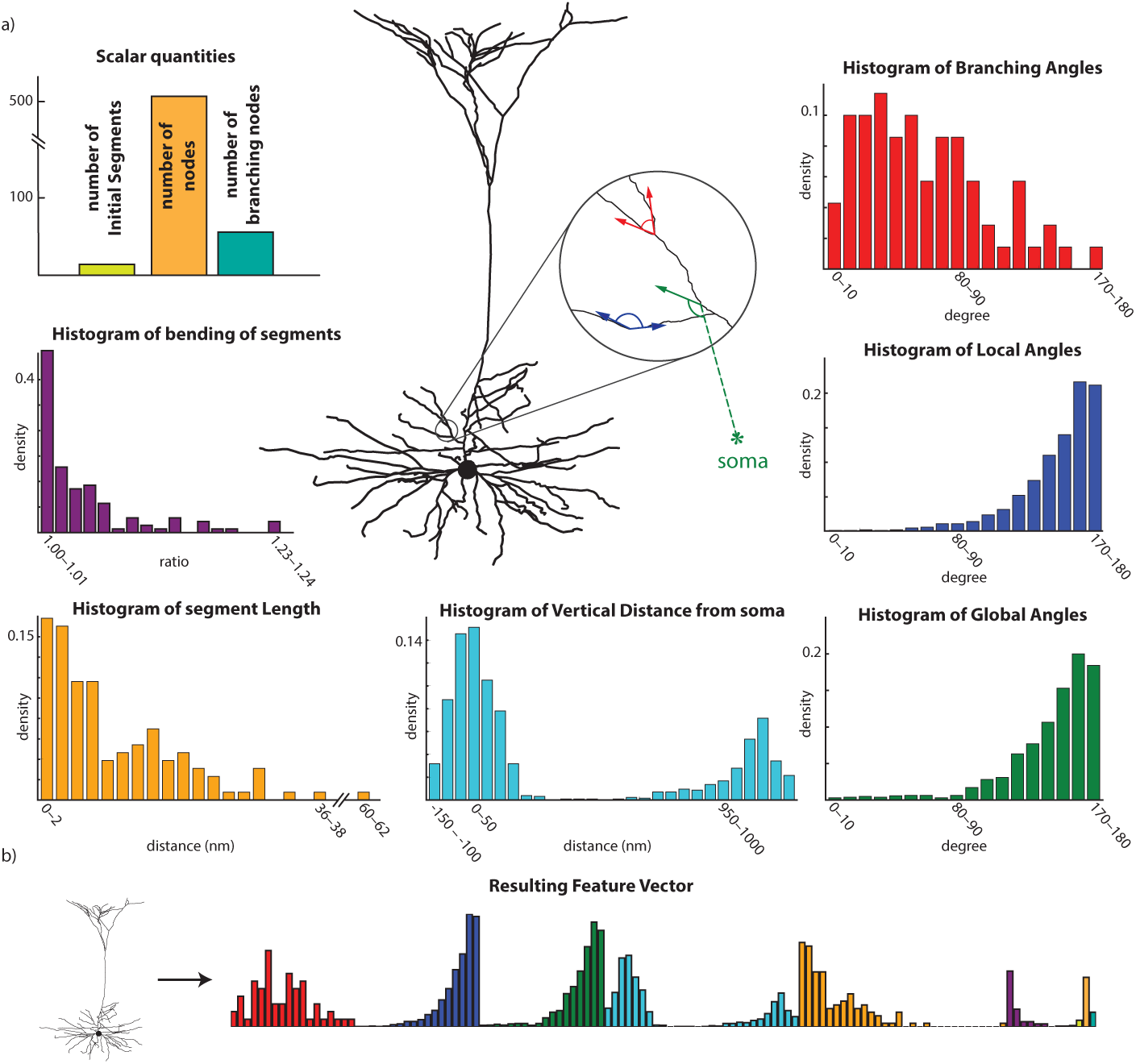
Representing a neuron morphology by its geometrical and morphological features. a) Morphology of a neuron is shown in the center, surrounding by its extracted features. b) By putting all the features together we can construt a long feature vector that describes is section 4.2

To attain a generative model for a neuron type, we want to be able to ask how representative a morphology is relative to those in a database. The morphologies in the database will all be different from one another because of randomness in the growth process, environmental conditions and, maybe disturbingly, details of the imaging techniques. To obtain a simple generative model we can characterize the database by the univariate means and variances of each feature (figure 2.a). To calculate the probability of a neuron under this generative model, we simply assume a normal model where each feature is assumed conditionally independent on the others, an assumption that is called naive Bayes. This gives a direct way of calculating the log probability of a morphology under the Gaussian model (figure 2.d) for each possible feature vector.

**Figure 2:**
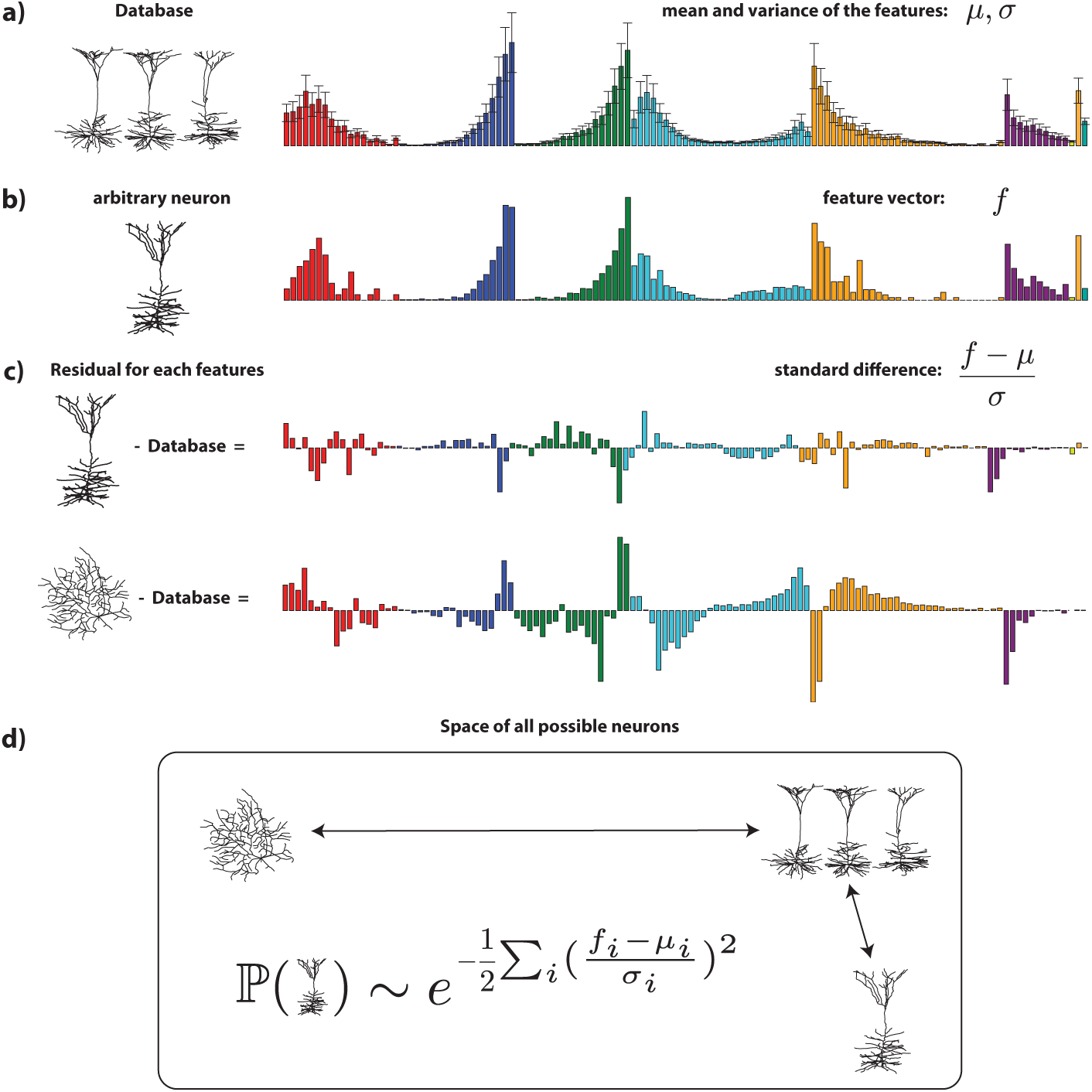
Features can be used to define a generative model of a given database of neuron: a) 3 samples of pyramidal neurons are shown on the left. On the right side the mean and the variance of features across all the samples are shown. b) An arbitrary neuron and its features are shown. c) By taking the standard difference between the arbitrary neuron and the database the residual of features for two neuron are computed. d) the square sum of all residuals (scaled by their inverse std) defines a metric for the distance from database. Moreover by taking the exponential of distance we can define a Gaussian probability distribution over the space of all possible neurons. (for details of generative model look at 4.3).

The number of possible neurons grows rapidly with the number of nodes. The degrees of freedom are three times the number of nodes. This makes it hard to know if the distribution of trees is meaningfully defined by the above defined distribution of features. One possible approach is to fix the dimension of the tree, e.g. by giving each neuron 1000 nodes. The other approach that we take in this paper is to correct the probabilities by normalizing them with a factor derived from the dimensionality of the tree (see Methods for details). With this assumption we can obtain a meaningful distribution over trees.

Once we have a generative model of trees based on feature similarity and dimensionality, we need a method for sampling morphologies. Here we used Monte Carlo Markov chain (MCMC) for sampling. MCMC is an efficient way to sample from a potentially nontrivial space. MCMC starts with an initial state and in each step, the current state will be perturbed to produce a proposal state (figure 3). We choose the Metropolis Hastings algorithm. *x* and *x′* are the current state and the proposal state, respectively, and ℙ(*x*|*y*) is the probability of jumping from state *x* to state *y*. The probability of acceptance of the proposal is equal to:

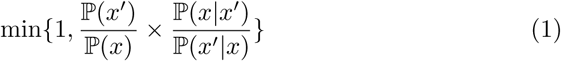

This allows sampling from a problem with fixed dimensionality.

To generate a diverse set of trees, the number of nodes needs to change during sampling. Reversible jump MCMC (RJMCMC) can deal with this problem by including the Jacobian matrix and correcting the acceptance probability ([29]). In this case the second term in the expression 1 should be multiplied with the determinant of the Jacobian of the mapping from the domain of the current state to the proposed state. By iterating this process, the features of the current neuron gradually becomes closer to those from the database during the so-called burn-in. It then subsequently samples morphologies from the generative model.

**Figure 3:**
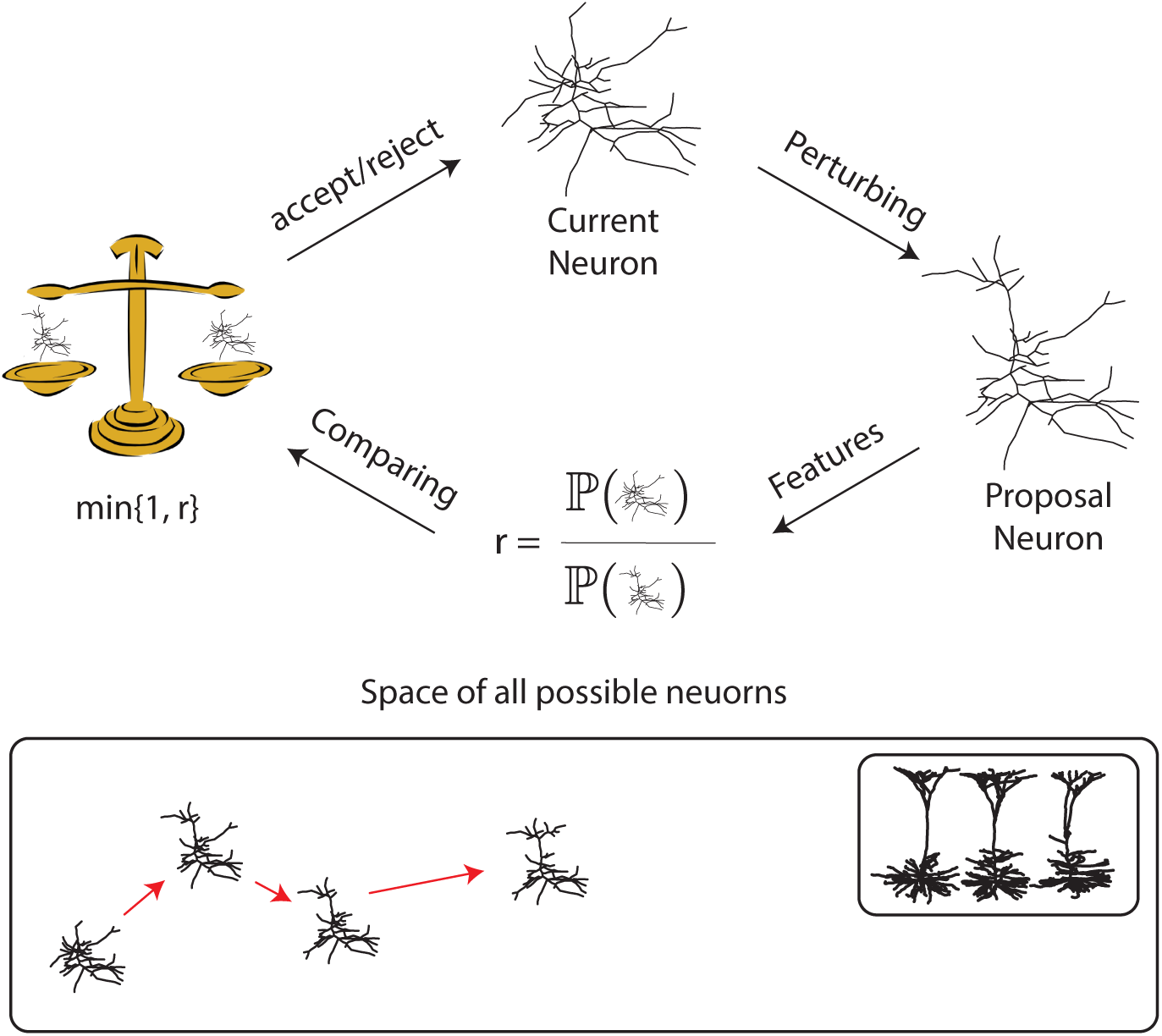
Schematic illustrating of MCMC algorithm for sampling neuron morphologies. When the morphology of a class of neuron is modeled by a probability distribution over the space of all possible morphologies, MCMC can sample. It starts with an initial neuron; possibly a trivial neuron (only soma) or an element of database. Iteratively, the current neuron is perturbed to make a proposal neuron. If the proposal neuron is closer to the database compare to the current neuron, it will replace it, otherwise it may be rejected. After many iterations the neuron gradually move toward the database.

Since the space of all possible neurons is huge and the features are different, mixing time, the time it takes for the chain to converge to the meaningful probability distribution, matters. To address mixing we tested different perturbation method to change the neuron in each iterations and selected those that allowed faster mixing. Our perturbations change the morphological structure (see figure 4) and doing so move the morphology towards the high probability region. Some of our steps propose to add or remove nodes. Since the number of nodes changes across iterations, this perturbation jumps between different dimensional space and hence we have to implement RJMCMC. Other steps propose rotations of part of the neuron. This facilitates convergence of the geometrical picture of the neuron in addition to the statistics of its segments. And yet further steps propose sliding parts. This enhances the convergence of morphological features. The combination of all of these proposals converges reasonably fast.

**Figure 4:**
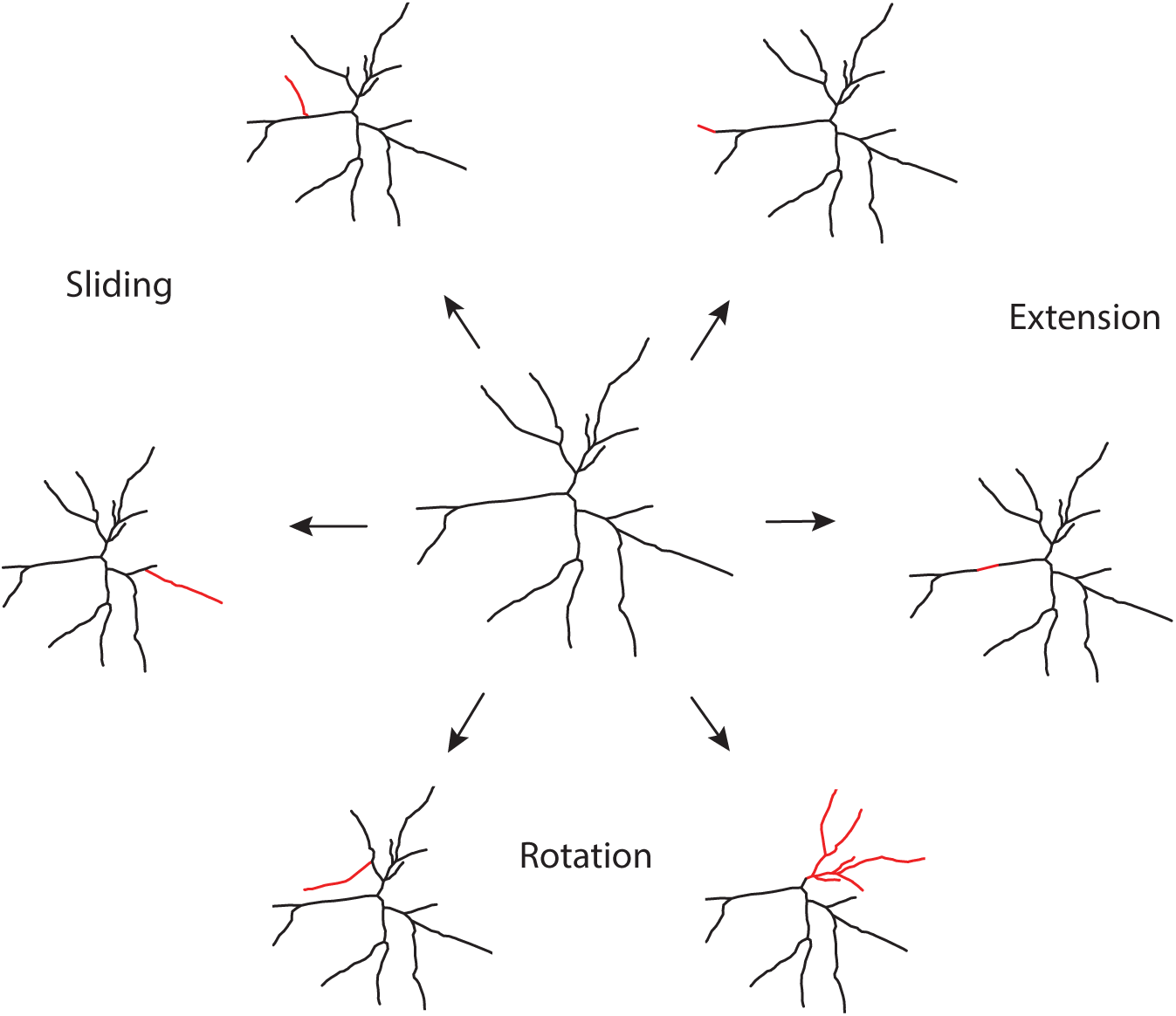
Proposal distributions. Some perturbation change the topological structure of the neuron and some of them change it geometrically (e.g. location). In the center an initial morphology is represented and around it are some possible proposals. These proposals usually change only parts of the morphology (red). In Sliding a part of the neuron moves over the morphology. This part is a connected component that attaches to random point on the neuron. In Extension a new node is added to an end point of the neuron. In Rotation one of the connected component of the morphology is rotated in 3d space.

### 2.2 Simple objects

To sanity-check our algorithm we first used it to generate simple objects. Since each neuron consists of segments we first test the algorithm to make a segment that is slightly curved. To measure this, we define bending as the fraction of the whole length of the object to the Euclidean distance between its endpoints. The goal is to generate segments with a bending ratio in the vicinity of 1.15. The algorithm starts with a straight line and in each step a random node of the segments will be selected and all the nodes on one side of it rotate. We find that the algorithm rapidly converges (Fig. 5.a). The algorithm thus seems to work well for simple problems of fixed dimensionality.

**Figure 5:**
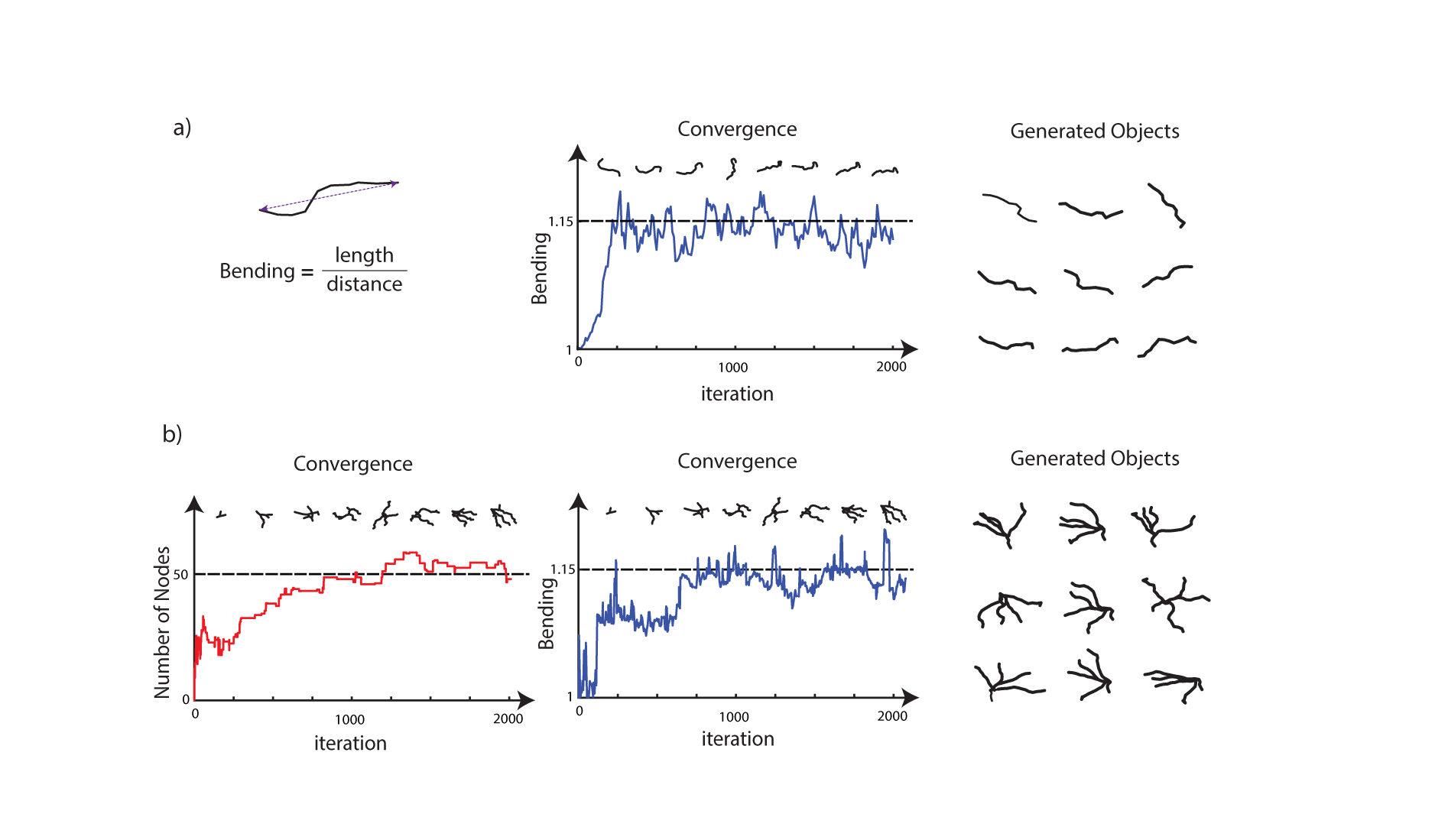
Generating simple morphologies. Neurons are assembled by connecting segments therefore generating them can be reduce to generating segments and trees. a) To generate a segment, we started with straight line and each time rotate a part of it using MCMC algorithm. The objection is to have the bending ratio close to *1.15*. The middle diagram shows the convergence during the iterations. Nine samples are plotted on the right. b) To generate trees with segments, we allowed changing the number of nodes during iterations. The objection here is to generate a geometrical graph with around 50 nodes and 4 branching nodes such that its segments have bending ratio around *1.15*. Convergence of number of nodes (red) and bending ratio (blue) for generating one object is plotted. Nine samples are plotted on the right.

To further check our algorithm we ran it on a case where the number of nodes does change. The objective is to both match the number of branching points, the number of nodes and also match the bending ratio. During this process the number of nodes can change. The algorithm readily achieves the defined objectives(figure 5.b). Asking for several features to be fit slows down convergence. Both the Metropolis Hastings and the RJMCMC parts by themselves seem to work well for simple, understandable, problems.

### 2.3 Generating Neurons

We now turn to our real objective, generating a morphology from a database of actual neuron morphologies. We start by analyzing layer 5 pyramidal cells in the neuromorpho database (6.a). Every neuron is different from one another but they share the morphological features of this neuron class.

To understand how our algorithm can replicate such a neuron, we analyze the evolution of the neuron over subsequent samples (6.b). We start with a trivial neuron, a soma with just seven equally spaced neurites. It takes the algorithm roughly 1000 samples to produce a morphology that roughly looks like a pyramidal neuron (upper). After another 25k iterations we obtain a morphology that looks quite real (lower right). Quantifying the features of the neuron reveals that some features converge quickly, e.g. distance (6.b upper) and others that converge quite slowly, e.g. the number of branching nodes (6.b lower). Importantly, when added across all features convergence happens, but very slowly. Indeed, even after 25k iterations they are probably not perfectly converged, which we share with most other real-world applications of MCMC. We thus see how the accumulation of small changes can give rise to complex tree morphologies.

We then test our algorithm on relatively distinct classes of neurons. Pyramidal, Tripolar, Purkinje and Stellate, in the database are considered (figure8 right). For each class there are about 1000 morphologies. As we see in the figure the generated morphologies look similar to the real ones (figure2 left). Some features are visually traceable in the generated neuron, like the density in different locations of space, yet other features are hard to inspect visually, e.g. the density of length of segments. Overall, the method works well across distinct classes of neuron morphologies.

**Figure 6:**
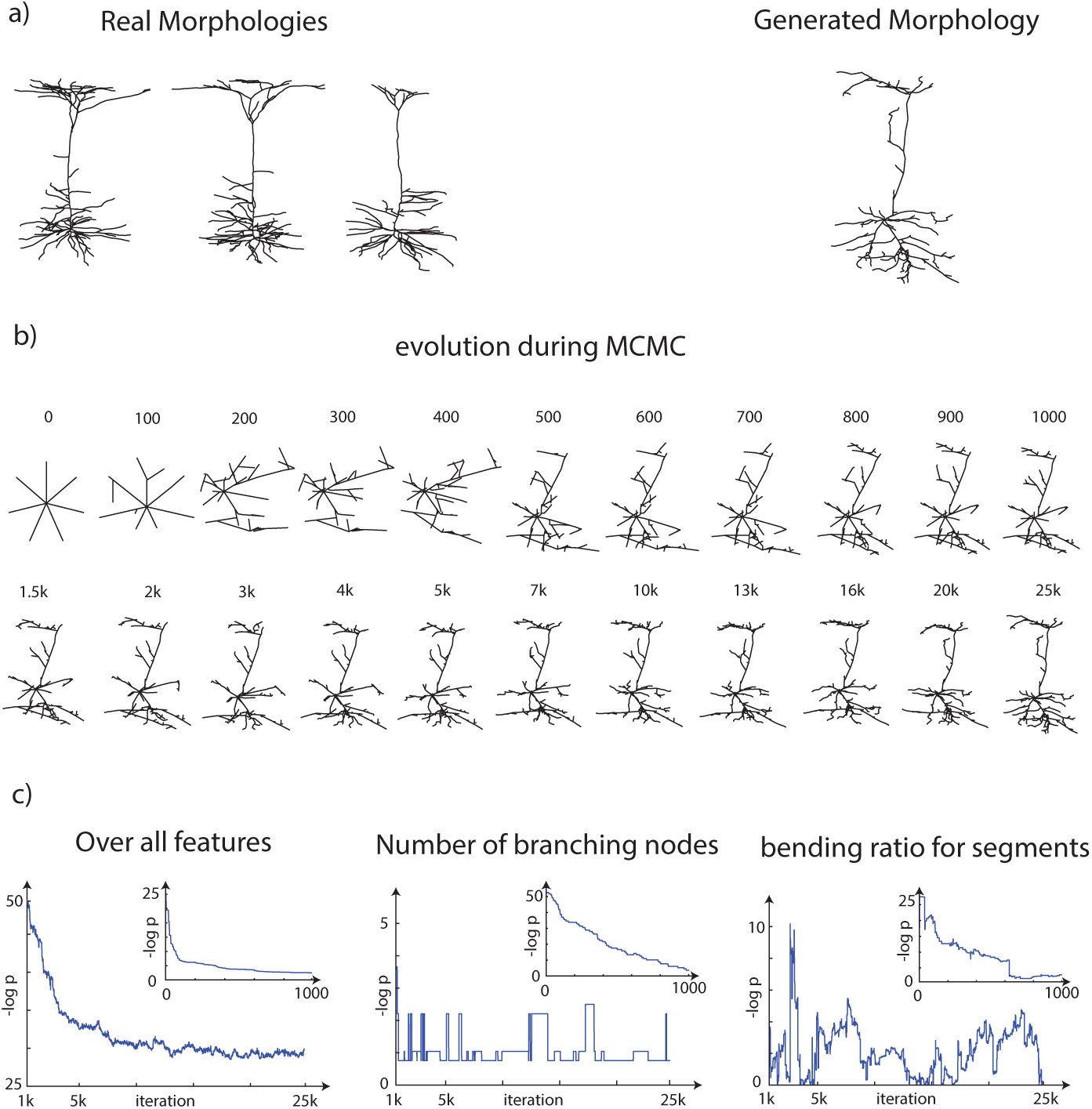
Generating a Pyramidal neuron: a) 3 real morphologies from a database of layer 5 pyramidal neuron are shown on left and a generated morphologies with our algorithm is shown on the right. The algorithm is run for 25k iterations and to depict how it converges a few samples during the iteration is plotted in (b). Above each neuron, the iteration's number is written. Since the general imagery of neuron did not change dramatically after 1000 iteration we plotted more neurons for the initial running. To explore the convergence we looked at the overall distance (left) as well a two features during the iterations (left and middle). The onset is the convergence up to 1000 iteration.

**Figure 7:**
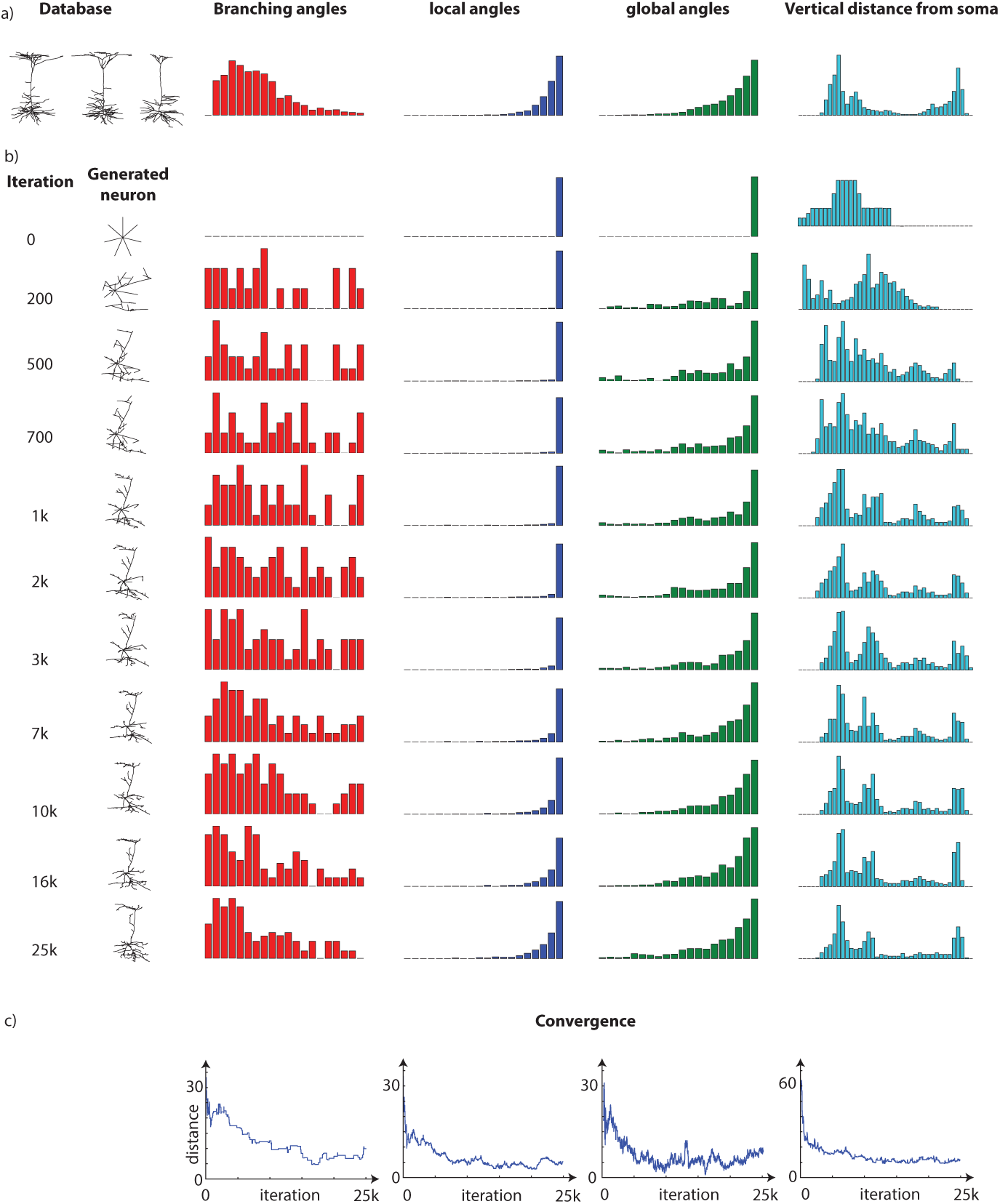
Convergence of features. We continue generating pyramidal neuron in figure 6 by plot four histogram during the iterations. a) the database and the mean of each feature are shown. b) the histogram are shown during the a few iteration number. c) the distance of feature from database if shown.

**Figure 8:**
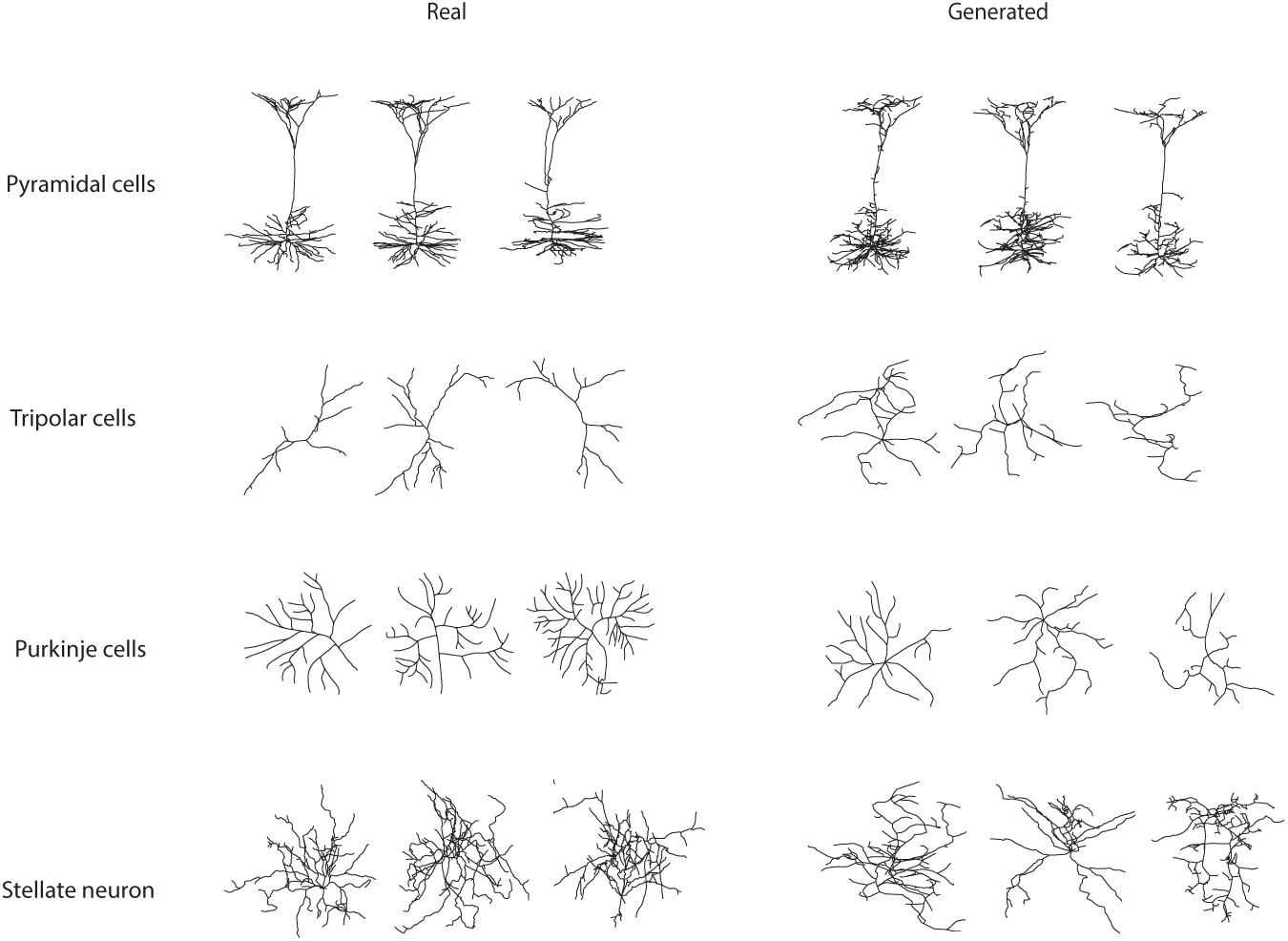
Generated neurons from each of four classes in the database. For each classes, three typical samples are plotted to the left and three generated, morphologies with our method are shown in the right. See supplement for additional information

### 2.4 Comparison with NeuGen

To conclude that our method is useful we need to compare to the current state of the art. We thus compare with the NeuGen package. NeuGen is a growth based model of generating morphologies that was developed in 2006 [19]. Although the generated neuron for both method looks good but when we look at a variety features RJMCMC shows its advantage (9). Moreover, it looks like RJMCMC produces a broader set of neurons. NeuGen also simulates the actual neural growth process, which is interesting for developmental applications, but our approach aims at just producing a matching morphology. RJMCMC produces morphologies that match the real ones in many ways and, in its focus on morphology quality vs process it offers a new tool to generate morphologies.

## 3 Discussion

We have introduced a method for generating dendrite morphologies. It extracts human-chosen morphological features from a database. It then uses the means and variances of these features to construct a simple generative model using the naive Bayes assumption. Sampling from the resulting distribution is challenging because morphologies may present with different number of nodes. Using the neuromorpho.org for dataset and RJMCMC for sampling, we test our algorithm and find that it meaningfully matches the statistical distribution of the training dataset and produces morphologies that look strikingly similar to us. We find that our method properly matches the full feature set where a traditional generator for morphologies only matched a subset of feature statistics.

Countless processes contribute to the generation of neuron morphologies but we only use a limited number of features in our algorithm. Regulation happens through countless molecular cascades [30], mechanical factors [31, 32], electrical activation [33] and geometrical limitations [34]. Each of these factors will be different, even across neurons of the same class, producing a highly structured multi-dimensional distribution[17]. And yet, neurons in one class typically share common characteristics, e.g. pyramidal neurons can be characterized by severally short basal dendrites and a large apical dendrite joined to an arborization in the tuft [35]. We have used a long feature vector (around 600 features). We can never be sure that there are no important features that we are missing. Future work, could use better features vectors to generate more meaningful morphologies.

An alternative to the human choice of features is to have algorithms choose those features. The recently popular framework of Generative Adversarial Networks (GANs) allows doing just that [28]. In that framework one network would figure out how realistic a neuron is by comparing the real and the simulated morphologies, while another system generates simulated morphologies. In a way, the system automatically finds the right features without human guidance. Using this method, GANs could successfully be trained to produce images of simple objects[36] or draw new paintings with the style of a given artist[37]. While exciting, it has been shown to be hard to generate a good statistical distribution using a GANs [38]. GANs may be a new hope for better generative models but they may suffer from statistical issues.

To convert our set of features into a generative model, we implicitly modeled them by an independent Gaussian distribution, which could not be further from the truth. First, we expect correlations of the features within a group, e.g. two close bins in the angular histogram (figure 1), or between groups; bending ratio (degree of straightness of the segments of a neuron) and the curvature. One could model the joint distribution in a more meaningful way, but that would be a hard statistical problem. Second, the assumption of Gaussian distribution might be violated for some features, e.g. those that represent counts. To overcome these issues, one can modify the generative model by using empirical distribution of the features from a big database. The generated morphologies from our method show that while assumption of independent distribution for features are far from real it can still produce satisfactory results.

While in theory RJMCMC samples evenly from the distribution, in practice we should expect it to have a large mixing time. Therefore, we should never expect chains to mix. To check that the neuron are meaningfully generated, we analyzed the convergence on the features presented (figure 6). However, this casual analysis cannot ensure that the chain properly mixes and generally there is no real solution to the convergence of MCMC[39]. Using a wide set of meaningful proposals enabled our technique to have a good acceptance rate and relatively quickly converge to high probability solutions. There are many ways how convergence behavior could be improved. For example, in the Ising model, the Wolff algorithm collectively flips the spin of a cluster of units and this can accelerate convergence by allowing the state to move non-locally [40].

Moreover, by using Hamiltonian MCMC we can potentially avoid random walk behavior in our sampling method [41]. Running multiple chains at different temperatures may also help improve mixing[42, 43]. Importantly, we do not claim that we evenly sample from the generative model but the fact that we are good at matching the features should be enough for many applications.

While the proposals are inspired by neuronal development, our method can not speak to the issue of developmental neurobiology. Our approach is fundamentally different from previous approaches which they use an explicit growth process [19, 20, 21, 22, 23, 24]. The biological inspiration of our proposals helped us speed up convergence. Moreover, the approach presented in this paper and previous growth-based works a may not be mutually exclusive. We could use their growth rules as part of our feature set. Or alternatively, we could initialize our algorithm with the results of their growth process, allowing a fine tuning of the results. Here we focused on de-novo generation of morphologies.

A good morphology generator could be useful for simulating a realistic neural network. After all, there is heterogeneity in morphology across neurons, and without a meaningful generator it is impossible to simulate networks of heterogeneous neurons. Sampled neurons could be used to find the link between the function of a neuron and its morphology [44] or its connectivity with other neurons [45]. With our methods its easy to generate samples of the morphology of any neuron type, which can facilitate realistic simulations.

One could also foresee that a good generative model could be useful for the segmentation of images. For example in electron macroscopy (EM) data a huge dataset of images is produced but there extracting the skeleton of each neuron is hard. Combining good generative models of neurons with current approaches promises better segmentations.

## 4 Materials and Methods

Neurons have a variety of morphological structures; Dendritic trees come in all shapes and sizes. They range from a total length of a few tens of micrometers to a few millimeters and vary significantly even within one neuronal class. Because of this wide spectrum of morphological shapes, building a growth model is usually difficult. On the other hand, the morphological structure can be characterized by a feature set. In this section, we start by a quantitative representation of neuron morphologies, a feature set. Based on these set, we construct a generative model and, using RJMCMC, sample from this generative model. This allows us to sample new dendritic trees that are statistically similar to those in the database.

### 4.1 Representation of a neuron’s morphology

The geometry of neuronal arborizations can be stored in swc format [46]. In this format the morphology of a neuron is modeled by geometric graph where the nodes and edges represent the point on the morphology and links between the points, respectively (figure4.1.a). For each node three pieces of information is provided: its three-dimensional location, the radius of the biggest sphere contained in the morphology and the neurite type at the location of the node (soma, axon and dendrite). Computationally, it is easier to extract the features of the morphologies and perturb them using this compact representation.

#### 4.1.1 Sub-sampling of Nodes

Some morphological features depend on the number of nodes in the swc file. To make them comparable across all neurons in a database, we use a subsampling method. Our method preserves the terminals and branching nodes and meanwhile selecting the nodes on a segment such that the distance between two consecutive nodes is within a bound. More precisely, it starts from one end node and greedy pick the maximum number of nodes that distance between the consecutive nodes is bigger than the threshold (figure 4.1.b). While the lower bound increases the number of nodes in the sub-sampling method decreases and the neuron is approximated poorer (figure 4.1.c). By choosing a fixed bound for the sub-sampling method, we can approximate all the neurons in a database with the same quality, which facilitates feature extraction.

### 4.2 Features Extraction

Graphical representation of the morphologies of a neuron alongside with the uniform sub-sampling of the nodes enables us to define their features. Some features we used are previously described as L-measure[47] and some of them are novel. Generally speaking, the features can be classified into 3 kinds: scalar, density and histogram. For the two later cases, we divide the whole range to bins and calculate the values in each bins. The type of each of the features and the size of feature (for the histogram the number of bin size) are written in the table 4.2. Here we shortly describe them:

**Table 1:** Features of the neuron: the list of all the features

| index | name of feature | type | method of quantification | size |
| --- | --- | --- | --- | --- |
| 1 | Number of nodes | scalar | value | 1 |
| 2 | Number of initial segments | scalar | value | 1 |
| 3 | Number of branching nodes | scalar | value | 1 |
| 4 | Number of end nodes | scalar | value | 1 |
| 5 | Global angles | vector | histogram | 20 |
| 6 | Local Angles | vector | histogram | 20 |
| 7 | Segmental branching angles | vector | histogram | 20 |
| 8 | Side branching angles | vector | histogram | 20 |
| 9 | Curvature | vector | histogram | 20 |
| 10 | Neural/Euclidean | vector | histogram | 20 |
| 11 | Segmental Neural/Euclidean | vector | histogram | 30 |
| 12 | Neural length of segments | vector | histogram | 50 |
| 13 | Self-avoidance | vector | values at meshes | 50 |
| 14 | Spatial distribution | vector | values at meshes | 50 |
| 15 | Fractal growth | vector | values at meshes | 10 |

#### 4.2.1 Number of nodes, branching, end nodes and initial segments

A neuron is built up by a binary tree except for the Soma. Four basic scalar features of this graph are the number of nodes, number of branching nodes, the number of end nodes and the number of nodes attached directly to the soma i.e. initial segments.

#### 4.2.2 Global Angles

Global angles measure how straight the segments of the neuron are grown away from the soma. It is computed for each node by the angle between the direction that neurite has grown with respect to position of the soma as the origin and are defined by measuring the vector connecting the node to its parent and the vector connecting it to the Soma. Since the neurons have the tendency to explore the surrounding space, it is expected that in many nodes this angle is obtuse.

#### 4.2.3 Local Angles

Local angles measure the straightness of the neurites by calculating the angles between two vectors: the vector connecting the node to its parent and the vector connecting the node to its child. Notice that the node has only one child to define the local angle. If the segment of the neuron is a flat line locally at the node, this value would be 180 degree. Lower values indicates the curvature of the neurite.

#### 4.2.4 Branching Angles

At each branching node, we can be calculate the branching angle by computing the angle between the vectors connecting the branching node to its children (figure 4.2.12). Usually this is an acute angle.

#### 4.2.5 Side Branching Angles

Since the neuron are 3d object, the branching angle itself can not describe the local shape of neuron around the branching node and we need to look at the side angles; the angle between the vector connecting a branching node to its children, and the angle connecting it to its parent(figure 4.2.12). Similar to the branching angle, we can approximate the side angle locally or segmentally.

#### 4.2.6 Curvature

Local angles describe the directional changing in the segments in each intermediate node, however these changes are correlated for close-by immediate nodes. The curvature measures between the difference in two consecutive local angles. In[48] the author used the same measure for analyzing the curvature of the segments of the neurons of drosophila.

#### 4.2.7 Spatial Distribution

Although neurons vary in the way they distribute in space, neurons from the same class usually share a similar spatial distribution. To quantify the spatial distribution, we rescale the neuron to put it in the unit 3d box. Then by meshing this box with a regular latices, the number of nodes that occupy a in every mesh can be counted. A schematic way of meshing is plotted in the figure 4.2.12. This feature is containing a rough approximation of Sholl analysis in particular[49] which first used to distinguish the visual and motor cortices of cats. In Sholl analysis the number of crossing for a circle of given radius is studied. We use a polar coordinate system to take into account symmetry orthogonal to the surface of many structures.

#### 4.2.8 Self-Avoidances

Neurites spread in space but usually avoid self-intersections. Similar to the spatial distribution we can measure self avoidances of the neurites by makeing a regular lattice and counting the number of nodes in each box. We count the number of boxes that only contains one node.

#### 4.2.9 Fractal growth

Similarly to the previous feature, we count the number of boxes that cover the neuron. This measure represent the fractal geometry of the the neuron [50].

#### 4.2.10 Length of Neural Segments

The length of each segment is an important property of a class of neurons.

#### 4.2.11 Bending

For each node the shortest neural path that connect it to the soma is usually close to a straight line. To make it concrete, for each node the ratio of its shortest path through the neuron to soma divided by the Euclidean distance between the node and Soma are calculated. By subtracting one and taking mean square of this ratio for all node we get the Neuronal/Euclidean ratio.

#### 4.2.12 Segmental bending

Similar to the previous feature, we can measure the flatness of each segment of neuron by dividing the neural distance and its Euclidean distance.

### 4.3 Generative model of the morphology

In this section, we define a probability distribution on the set of all possible morphologies given a database. In the previous section we defined a set of features for one neuron. We calculate the mean (*μ*_*i*_) and variance (*σ*_*i*_) of the features. Here we want to define a probability distribution over the space of all possible neurons such it concentrates around the points with the features of closing to those of the database. To fix translation invariant, we set the location of root (soma) to be at the origin. Suppose a neuron is made of *n* nodes and 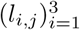 is the vector in ℝ^3^ that connect jth node to its parent (1 ≤ *j* ≤ *n* – 1). Also, *a*_*i*_, is the *i*th feature of the neuron. The probability distribution of the neuron is then proportional to:

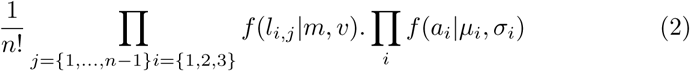

Where *m*, *v* are two predefined values and:

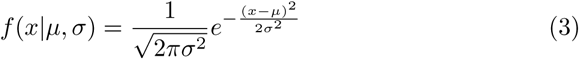

Notice that the probability distribution is made of two products; the first factor is independent of the features and is defined such that if no feature is defined, then integration of equation 2 over all possible neuron is finite (and is equal to one) and therefore the proposed probability distribution is well-defined. Keep in mind that once the length *l*_*i*_’s are fixed, then there are *n*! possibilities for connecting them to make a graph of tree. The second factor ensures that a neuron with the features close to the database has high probability value and therefore it has higher chance to be sampled.

Finally we have to define *σ*_*i*_’s. Naively, we would like to choose it to be the standard deviation of feature *i* in the database. However, if we do so, we obtain a rather bizarre problem. The effective measure of models is not independent of the *σ*_*i*_ for two reasons. First, features are not actually independent of one another which could be ameliorated by introducing extra parameters. Second, the number of possible trees co-varies with the *σ*_*i*_. There could be potential mathematically beautiful solutions to this problem. However, here we chose a simple pragmatic solution. We multiplied each *σ*_*i*_ with 5 and each histogram feature we also divided by the number of bins. We find that this ad-hoc strategy reasonably corrects for the two mentioned biases.

#### 4.3.1 Markov chain Monte Carlo

To use the generative model it is necessary to have a way to draw samples form the distribution. Here we use Markov chain Mote Carlo method to generate samples of neuron morphology. It start from an initial neuron and in each iteration one of the perturbations is selected randomly (the perturbations are explained in the next section) to obtain a proposal morphology. In Metropolis Hasting, if the probability value of the proposal neuron is higher, it will replace the current neuron otherwise with the probability of their ratio (proposal to ratio) the proposal will be accepted. When the Markov chain is not symmetric, the acceptance probability should be modified to:

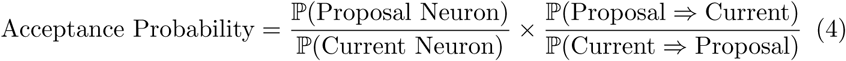

or in other words:

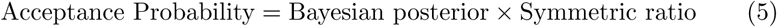

The first ratio indeed compare two states and is called Bayesian posterior. The second ratio, *symmetric ratio*, is added to make the Markov chain symmetric and it should be calculated for each perturbation individually. For many of perturbation this ratio is 1 but especially for the perturbation that changes the dimension of the neuron (increase or decrease the number of compartments) it should be elaborately calculated.

#### 4.3.2 Reversible jump Markov chain Monte Carlo

When the dimension of current and proposal neuron are different, neither of the ratios in equation 5 are well defined. Reversible jump Markov chain Monte Carlo is a way to get around this problem by replacing the first ratio with the ratio of the probability density function of two probabilities and the second ratio with the Jacobian of the mapping between the two space([29] and [27]). In the list of proposals below, the first one needs this modification and we calculate and simplify the ratio there.

#### 4.3.3 Initialization

An initial neuron is required to run the algorithm. We need a simple neuron with one node as soma and a chain of connecting nodes. For certain questions, generating neurons de-novo is not necessary and initializing with real neurons could leads to better results. To validate the method, we tried both ways of initialization.

### 4.4 list of proposals

To preform MCMC on neurons, we need a set of actions to perturb the neurons. When they act on a neuron, the shape of a neuron changes and therefore the features of the resulted neuron are different. Based on the probability density of the two neurons and the RJMCMC factor, the new neuron may be replaced with the initial one. Here we present the list of all perturbations on a neuron. Some of the perturbations change the neuron geometrically, for example rotating a part of neuron, some of them also change the graphical structure of neuron, for example by perturbing the connectivity between nodes. The perturbations used in this paper can be classified into three categories.

#### 4.4.1 Extending or Reducing the neuron

The neuron is made of a set of nodes and in this perturbation a node is added or removed. There are different ways to do this perturbation. When one of these sub perturbations are chosen, the node in selected uniformly from all possible nodes. The probability of selecting one of these perturbations and the possible way of choosing one node is shown schematically in the figure4.2.12.a. When the new node is created its location is drawn from a 3 dimensional Gaussian (with *m* and *v* as the mean and variance). When the created/removed node is an intermediate node, all the nodes after the created node are shifted by the location of the new node. Notice that the jumping from one space to another is made by projection onto multiple Euclidean spaces and therefore the RJMCMC ratio for each of these perturbations are computed based on the number of possibilities and the reverse perturbations; and their ratio is generally not one. We define the number of possibilities of each neuron to be the number of potential jumps in figure4.2.12.a:

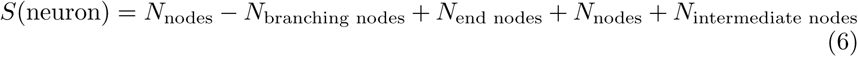

Then by using the equation 2, the probability of acceptance for extension is equal to:

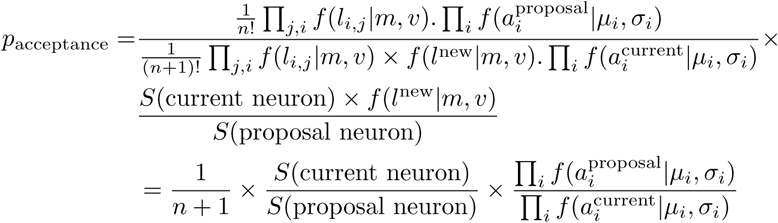

Where *l*^new^is the new vector added to the current neuron. Hence for example if the perturbation “Removing one end-node from the set of nodes” is selected, the RJMCMC ratio is equal to number of possibility for proposal neuron divided by the number of possibility for the current neuron.

#### 4.4.2 Sliding a part of the neuron over itself

In this perturbation, the neuron is detached from a node to turn to two separate parts, then these two parts parallel transport and reattach in the another node of the neuron(figure4.4.b). As the result the topology of the neuron changes. The detached node can be any non-soma node and the reattached node should be a non-branch node. These two nodes can randomly be chosen from all the possible nodes of the neuron but to boost the MCMC, we used two ideas. First, the distance between the these two nodes are less than a certain limit. Second, in many neuron classes it gives a better result when the neuron is detached from one of the two outgoing segments of a branching node of the morphology. Because of that we put a different probability for choosing the detached parts of the morphology among the branching nodes verses non-branching nodes. Notice that this perturbation is symmetric and hence the relevant ratio is one.

#### 4.4.3 Rotation

In this perturbation a part of neuron will be rotated along one node (figure 4.4.c). Specifically, a nodes will be selected and the part of the connected component which is not containing the soma would be rotated with a random unitary matrix in 3d. Similar to the previous perturbation, to boost MCMC we put different probability for selecting a uniformly random nodes of a neuron or a branching node.

The unitary random 3d matrix should be selected randomly from all unitary matrices. For boosting MCMC, it is better that the selected rotation be close to identity matrix and the probability distribution on all unitary matrix being symmetric. For doing that, we put a symmetric distribution on the space of 3d rotations which concentrate mostly around the identity. Since every rotation can be expressed as

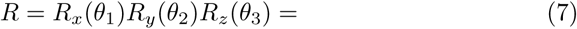

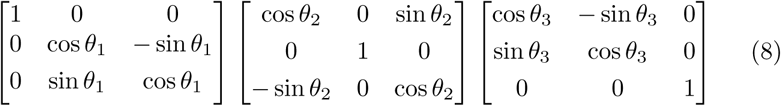

we can set a probability distribution on the *θ*_*i*_’s and hence make a probability distribution on the space of 3d rotations. The probability distributions on the *θi’s* are coming from one of the Von Mieses distributions which are symmetric distribution on the one dimensional circle. To symmetrize the 3d rotation, we use a simple trick of multiplying them in the form of:

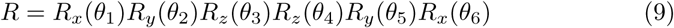

Notice that by doing that the probability density at the matrix *R* is the same as *R*^-1^.

### 4.5 Ergodicity

At the end, it should be taken into account that MCMC works only when the Markov chain is transitive, and here it can be checked easily it is the case. For example by removing the nodes from a given neuron we can produce a single node neuron (with *p* > 0) and by adding nodes we can go from single-node neuron to any neuron (again with *p* > 0).

### 4.6 Implementation Details

The algorithm implemented in python with three main classes. Neuron class gives a neuron object containing nodes. Each node has geometrical attributes (location and type) in addition to topological ones (parent and children). Moreover it calculates the features indicated above once the object is called. The second class take a database of neurons and extracts the parameters for the algorithm. The third class does MCMC sampling starting from a simple neuron and by applying the proposals perturbations on the initial neuron in each iteration shapes the neuron closer to the database. The code is available at: Neuron Generator

## 5 Acknowledgments

The authors thank Pavan Ramkumar and Hugo Fernandes for their significant discussion. Roozbeh Farhoodi was supported by Cognitive Sciences and Technologies Council (COGC) and National Foundation of Elites of Iran. Konrad Kording and Roozbeh Farhoodi were supported by NIH (U01MH109100, R01MH103910). KK initiated the first idea. KK and RF the mathematical foundations of idea. RF developed the software. KK and RF wrote the manuscript.

## 6 Supporting Information

Here we will give more details on the generation of various aspects of the simulations. Figure 1: The neuron was chosen from the Chen contribution (pyramidal neocortex) with neuromorpho id of 32114. It was then sub-sampled such that the distance between two consecutive nodes is around 20*μm*. The histogram figures are normalized such that the summation of densities over all the bins is equal to one. For each histogram related to the angles we chose 18 bins. For others the bin length is selected such that the number of bins is between 20 to 30 (see code-base for details).

Figure 2: 3 samples from Chen contribution (pyramidal neocortex) is shown. The mean and deviation is computed over the whole database. The features that presented here are the same as the feature in fig 1.

Figure 5: The objective function is 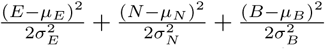 where *E*, *N* and *B* are bending ratio for segment (see the method section for definition), number of nodes and number of branching, respectively, and *μ*_*x*_ and *σ*_*x*_ are constant values representing the mean and deviation of the feature *x* (*μ*_*E*_ = 1.15, *σ*_*E*_ = 0.05, *μ*_*N*_ = 50, *σ*_*N*_ = 5, *μ*_*B*_ = 4, *σ*_*B*_ = 0.5). The proposal for first simulation (a) is the rotation around random node and for the second simulation (b) is the general sliding and the rotation around random node (see the method section for definition of proposals). The initial state for the first simulation (a) is a straight line made of 50 nodes with equal distant from parent. The initial state for the first simulation (a) is one node. Both simulations run for 2000 iterations.

Figure 6 and 7: pyramidal neocortex from Chen’s lab was selected and subsampled such that the distance between two consecutive nodes is around 20μm. The initial neuron of the algorithm is a 2D star shape neuron with 7 wings and 70 nodes on each wing and one node for soma. The probably distribution for selecting the preturbations is: ‘rotation for any node’: 3/13, ‘rotation for branching’: 4/13, ‘sliding general’: .5/13, ‘sliding certain in distance’: 1/13, ‘sliding for branching node’: .5/13, ‘sliding for branching node certain distance’: 1/13, ‘sliding for end nodes’: 2/13. The kappa for rotations are set to 400. the interval for sliding has the length 100. The MCMC is run for 25000 iterations.

Figure 8: Four different class of neuron morphology are selected: 1) Pyramidal cells of rat hippocampus from Chen database, 2) Tripolar cell of rat neocortex from Brown archive, 3) Purkinje cells of mouse cerebellum from Kengaku archive and 4) Stellate cells of mouse neocortex from Ballester-Rosado archive. For each database the features are extracted based on the table 4.2. The initial neurons are star-like neuron with 7 wings. all the samples run for 25k iteration. The detail of selecting perturbation is the same as 6.

Figure 9: The neurons that generated from NeuGen are pyramidal L5. To generate them, we changed the seed number to a range of different values. The output of the software are .hoc files. We converted them it to .swc file by finding the location of the mean of each component in the *.hoc file.

**Figure 9:**
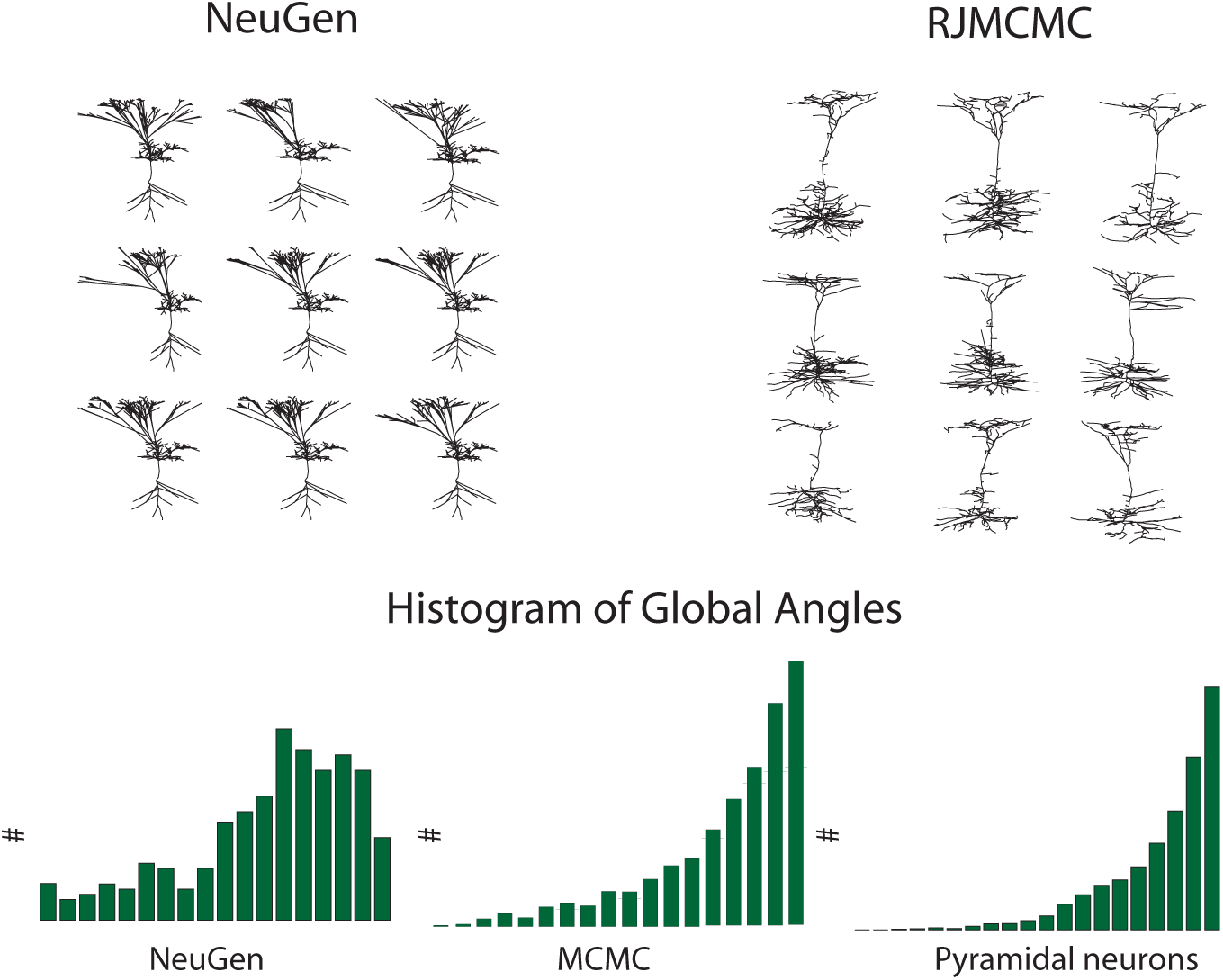
Comparing with growth based algorithm. We compare the algorithm that presented here with a state-of-art algorithm, NeuGen. It is designed for specific classes of neurons and among them we generated layer 5 pyramidal neurons. a few samples is shown on top left. A few samples from our algorithm is shown on right. In middle a histogram of global angle is shown for NeuGen, out method and Pyramidal cells. Global angles is one of the parameters that is not explicitly used in the growth process of NeuGen and the figure shows that the generated neuron failed to have it as their histogram is far from pyramidal cell. One of the aspect of our method compare to other package (including NeuGen) is that is fits better on the features and also it gives a degree of freedom to the generated samples. On bottom we chose a few features and extract the mean and variance from generated samples (for the angles, the mean over bins is computed). The error bar shows that the samples generated by our method have higher degree of freedom.

**Figure 10:**
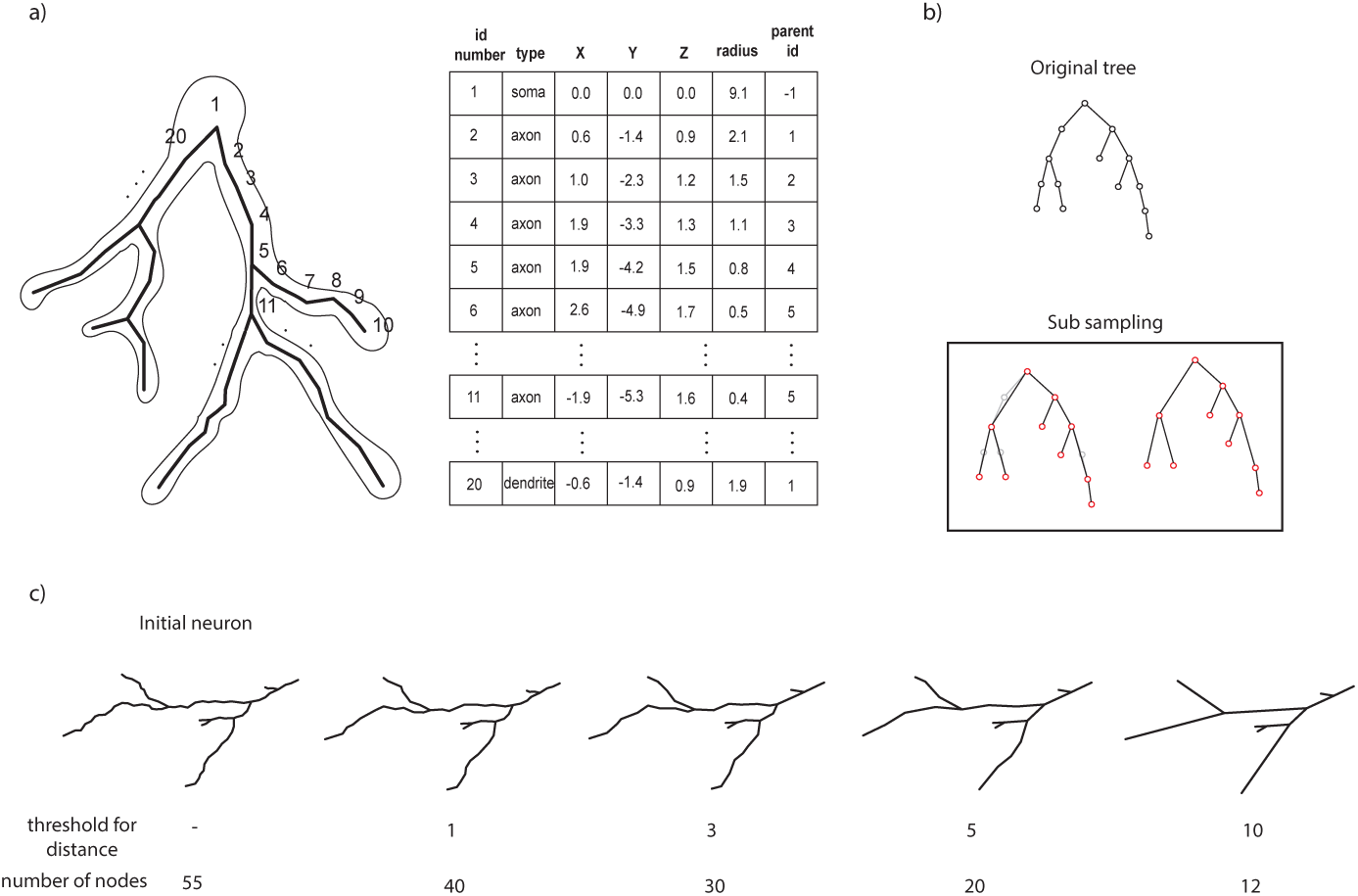
Representation of a neuron. a) neurons are constructed by a set of cylinders and spheres which have locations and diameters with an underlying tree structure (swc format). b) In this representation the distances between two consecutive nodes is arbitrary. However, for calculating many features of neuron, these distances should be roughly equal. As such we do sub sampling the neuron to straighten it. In this process on each segments of the neuron, the nodes that are closer to each other less that a certain threshold will be removed. c) as the threshold for distance increases, the sub sampling approximation become coarser.

**Figure 11:**
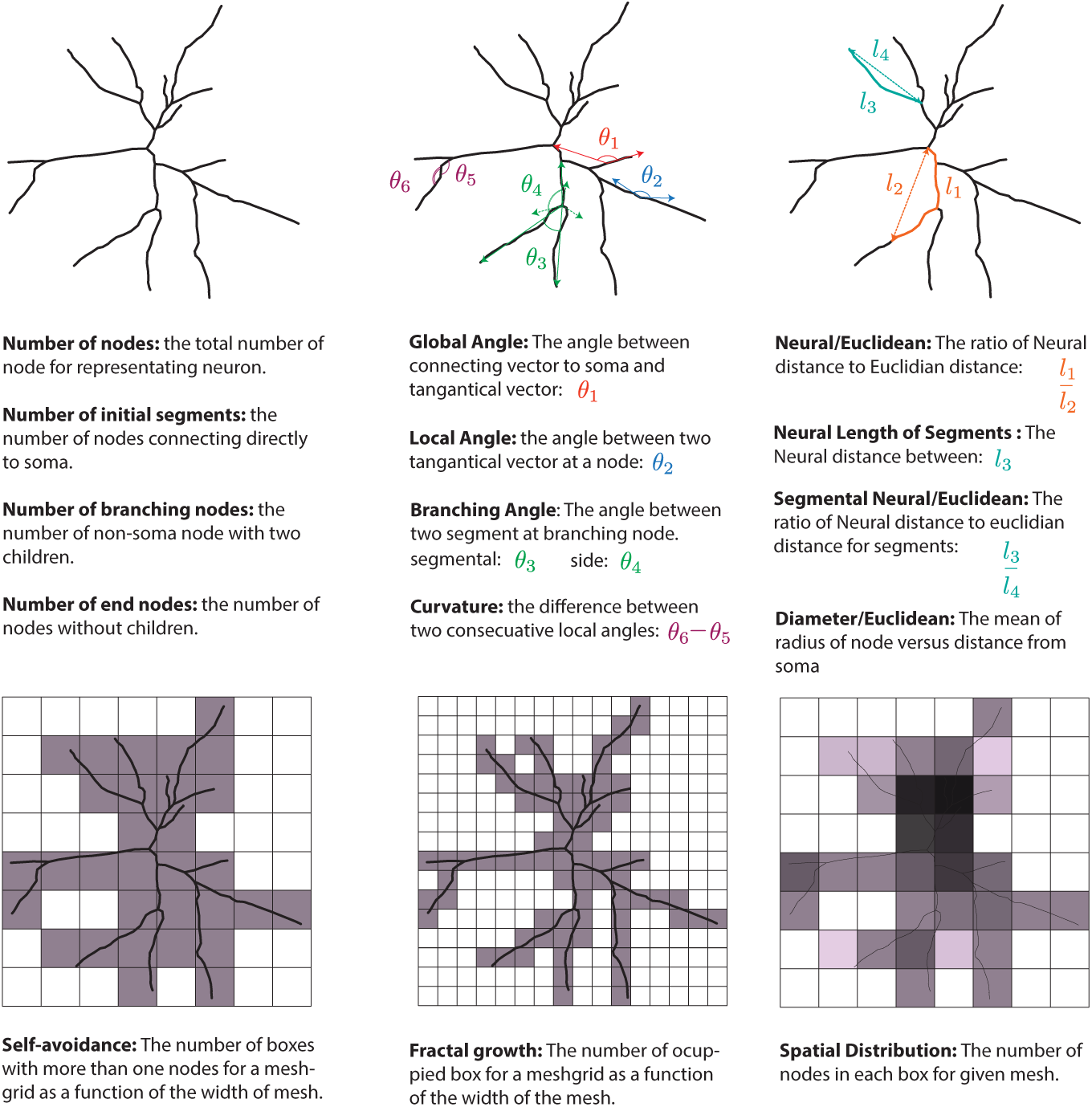
Features of neuron.

**Figure 12:**
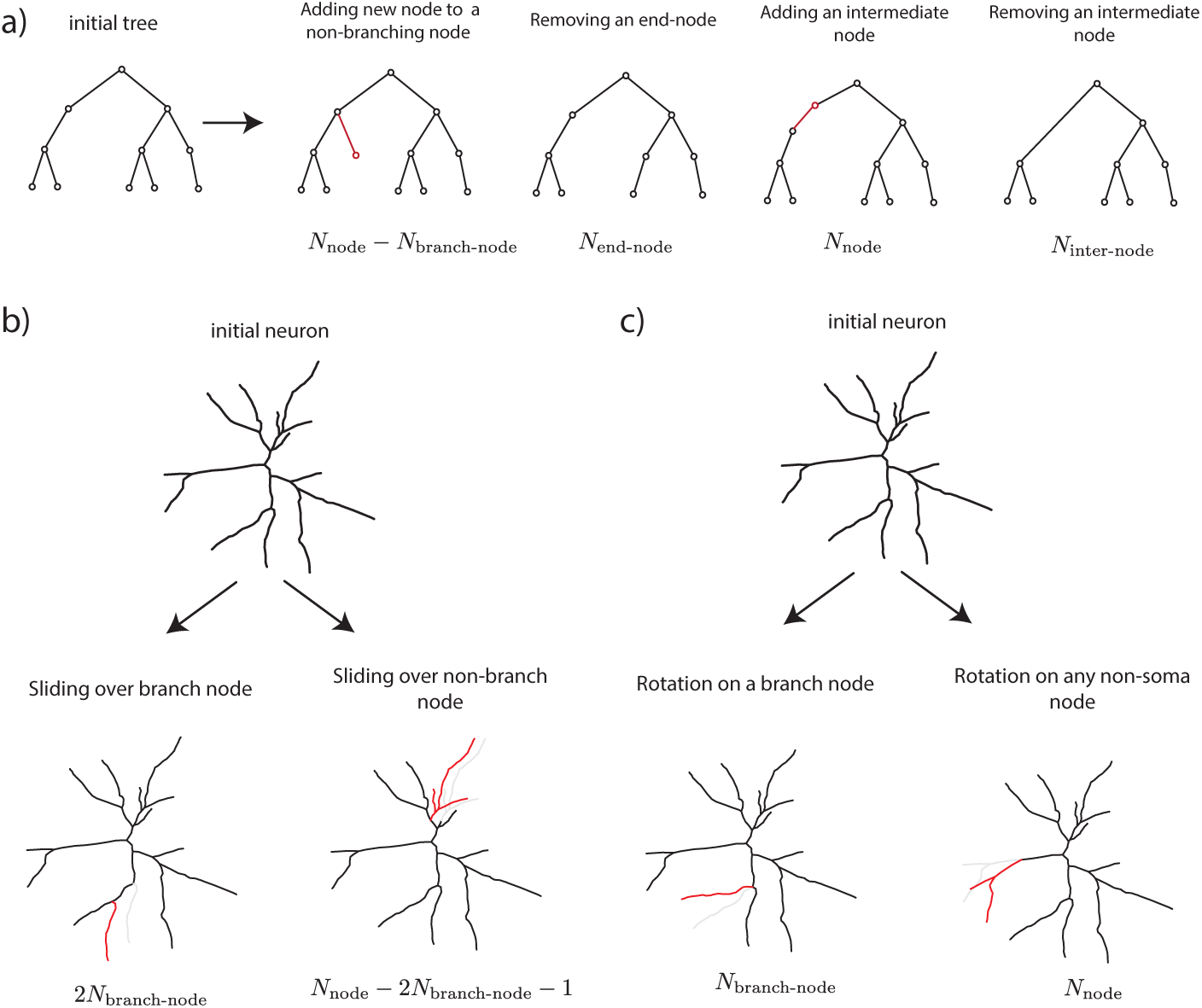
The list of all proposals a) flowchart of Extension/Reduction of a neuron. Starting from an initial tree, here we used two forms of perturbations. On the left side, an end node is added or removed while on the right side, an intermediate node is added or removed. The new or removed nodes are chosen randomly among all the possibilities which is written on the last line. The notations: N_end_ number of end nodes, N_inter_ intermediate nodes and N_branch_ number of branching nodes.Hence the number of all nodes of the neuron is equal to: N_node_ = 1 + N_inter_ + N_end_ + N_end_. Notice that there is not any limitation on the number of nodes that attach to the soma, but other nodes can at most have 2 children. b) The flowchart of sliding perturbation of a neuron. To slide the neuron over itself we need to select a non-soma node for detaching and nonbranch node for reattaching. When the non-soma node is selected, neuron is detached from the parent of this node and this two sections parallel transport and would be reattached in non-branch node. In the figure the gray part is the old position of one of the section and red one is the new position.The non-soma node can be the children of a branching node (left neuron) or any other non-soma node (right). Also the sliding distance (I in the figure) is forced to be less than a threshold. c) rotation along a random node. To rotate a part of neuron we have to select a non-soma node and rotate the part of neuron which is connected to it and does not contain the soma. The unitary matrix for rotation is coming from a symmetric distribution on the set of all unitary matrix. The selected node for rotation can be a node in general (left) or branching node (right)

